# HSF4: A Prognostic Predictor and Potential Therapeutic Target in Colorectal Cancer with Implications for Patient Immune Status

**DOI:** 10.1101/2025.06.25.661652

**Authors:** Ting Wan, Jun Wu, Wei Chang, Yahui Li, Can Shen, Jinghua Liu

## Abstract

Colorectal cancer (CRC) is among the most prevalent malignancies globally, characterized by high morbidity and mortality rates. The Heat Shock Factor (HSF) family of genes plays a critical role in the cellular response to environmental stress, particularly heat stress. These genes encode transcription factors that regulate the expression of heat shock proteins, which are essential for maintaining cellular homeostasis under stressful conditions. Recently, there has been a growing interest in the role of HSF family genes in the development and progression of CRC. In this study, we conducted a comprehensive analysis of CRC samples from The Cancer Genome Atlas (TCGA) database to identify potential therapeutic targets. A total of 533 gene expression matrices, were downloaded and processed, including 42 normal and 491 tumor samples. Differential expression analysis of the HSF family genes, revealed HSF4 as the most significantly differentially expressed gene between normal and tumor samples. Consequently an HSF4-related signature was constructed, stratifying all CRC patients in the TCGA dataset into low-risk or high-risk groups based on HSF4 expression. Survival analysis demonstrated a significant association between HSF4 expression and clinical outcomes indicating that the signature possesses substantial. Correlation analysis further revealed significant relationships between HSF4 expression and clinical features including Stage, T stage, M stage, and N stage. Univariate and multivariate Cox regression analyses confirmed the prognostic significance of HSF4. Functional analysis of differentially expressed genes based on HSF4 expression levels demonstrated enrichment in critical biological processes and pathways. Furthermore, HSF4 exhibited unique properties within the tumor microenvironment (TME). Additional analysis indicated that different risk groups exhibit varying levels of immune cell infiltration and cell proliferation capacity. Notably, the low-risk group showed increased sensitivity to multiple chemotherapeutic agents due to its lower IC50 values. In conclusion, this study presents the first signature for predicting the prognosis and immunological status of CRC patients based on HSF family genes. Our findings suggest that HSF4 may serve as a potential therapeutic target in CRC.

## BACKGROUND

Colorectal cancer (CRC) is the second leading cause of cancer-related mortality worldwide, possing a significant health challenge due to the projected increase in its incidence and mortality in the coming decades (Sawicki et al., 2021). Global health statistics, indicate that the burden of CRC is particularly high in developed regions largely attributable to dietary and lifestyle factors. However, developing countries are also experiencing a rapid rise in both cases and fatalities. This trend is driven by urbanization, dietary changes and increased life expectancy, which collectively contribute to the growing prevalence of the disease (Hossain et al., 2022).

The therapeutic landscape for colorectal cancer (CRC) has undergone a profound evolution over the past few decades. Significant advances in surgical techniques, chemotherapeutic agents, and targeted therapies have led to notable improvement in the outcomes of early-stage disease(Benson et al., 2021). Despite these advances, CRC remains highly treatable when detected early, with surgical resection and adjuvant therapies offering the potential for a cure. However, the high recurrence rate and the increasing incidence of drug resistance in advanced stages underscore the urgent need for the development of novel therapeutic targets and strategies to prevent disease progression and recurrence(Yang et al., 2024).

Among the various molecular players implicated in CRC, the Heat Shock Factor (HSF) family of transcription factors represents a promising avenue for research and therapeutic intervention(Dastidar et al., 2023; Gomez-Pastor et al., 2018). These factors are primarily known for their role in cellular stress responses, particularly in mediating the heat shock response, which facilitates cellular protection against stress by promoting the correct folding of proteins and preventing of aggregation of stress-denatured proteins (Akerfelt et al., 2010). In addition to this well-established role, HSFs are increasingly recognized for their contributions to cancer development and progression influencing cell proliferation, differentiation, and survival.

In cancers, the dysregulated expression and activity of HSF4 may contribute to the maintenance of proteostasis under conditions of cellular stress induced by genetic alterations and environmental factors. Furthermore, HSF4 may promote the expression of chaperones and other proteins that are essential for cancer cell survival and proliferation(Puustinen & Sistonen, 2020).

Given the intricate network of molecular pathways orchestrated by HSFs in cancer, targeting HSF4 presents a promising therapeutic strategy to disrupt proteostasis, inhibit cancer cell survival, and potentially overcome drug resistance. Further research is necessary to elucidate the specific roles of HSF4 across various cancer types and its interactions with other oncogenic pathways, in order to fully realize its therapeutic potential in cancer treatment.

This study aims to elucidate the role of HSF4 in the progression and prognosis of colorectal cancer by utilizing a robust dataset from The Cancer Genome Atlas (TCGA). This dataset consists of 533 individual samples from patients diagnosed with colon and rectal adenocarcinomas (COAD and READ), encompassing both cancerous and normal tissue samples, thereby serving as a valuable resource for in-depth genetic and transcriptomic analysis. Through comprehensive data analysis, which includes differential expression studies, survival analyses, and correlation assessments with clinical features, this research seeks to define the impact of HSF4 expression levels on disease outcomes and its potential utility as a prognostic biomarker.

This study’s further investigations employ advanced statistical and bioinformatics techniques to analyze the relationship between HSF4 expression and various clinical parameters including tumor stage, metastasis, and patient survival rates. Additionally, functional enrichment analyses and drug sensitivity assessments are conducted to explore the biological pathways associated with HSF4 and to identify potential therapeutic targets that could be leveraged for the development of new treatments for CRC.

This research not only enhances our understanding of the molecular underpinnings of colorectal cancer but also contributes to the broader field of cancer genomics by emphasizing the potential of transcription factors such as HSF4 as biomarkers and therapeutic targets in the clinical management of cancer.

## MATERIALS AND METHODS

### Publicly Data Collection

We downloaded gene expression data and clinical information for 533 samples from the TCGA database (https://portal.gdc.cancer.gov/), specifically for TCGA-COAD and TCGA-READ. This dataset includes 42 normal samples and 491 tumor samples. The expression matrices were extracted and merged using Strawberry Perl version 5.30.0.1.

### Identification of differentially expressed genes (DEGs)

Initially, we applied pre-filtering approaches (including the elimination of the genes with ‘all zeros’, and the removal of ‘NA’ values. Subsequently, we performed gene-wise standardization. Following this, we utilized Voom normalization and the Limma R tool were then utilized consecutively to identify the differentially expressed genes, with Limma employing the Wilcox test. This process resulted in the identification of a set of statistically significant genes. We then generated a volcano plot using bi-filtering approaches, which included p-value filtering and fold change filtering. A up-regulated gene is defined as one with a p-value less than 0.05 and fold change greater than 2, whereas a down-regulated gene is defined as one with a p-value less than 0.05 and fold change less than 0.5.

### The risk signature generation in CRC

Based on the median expression level of the most significantly differentially expressed HSF family gene in tumor and normal tissues, we classified patients into high-risk and low-risk groups. Subsequently, we conducted univariate and multivariate Cox regression analysis to confirm the association between this risk signature and patient prognosis.

### Pathway and Function Enrichment Analysis

After obtaining DEGs using the limma package, we proceed to conduct enrichment analysis utilizing both the Kyoto Encyclopedia of Genes and Genomes (KEGG) and Gene Ontology (GO) databases. This process involves annotating the genes with their corresponding biological functions and pathways, thereby enabling us to gain insights into the underlying biological mechanisms associated with the observed changes in gene expression.

### Cell proliferation scoring

To investigate the relationship between HSF4 and cell proliferation, the gene set “CELL PROLIFERATION GO 0008283” was downloaded from the GSEA database (https://www.gsea-msigdb.org/). Single sample GSEA (ssGSEA) analysis was employed to quantify the cell proliferation capacity of each sample, and a correlation scatter plot was generated using the ggplot2 package.

### Analysis of Immune Cell Infiltration and TME

To characterize the immune landscape of CRC patients, ssGSEA analysis were used to quantify the infiltration abundance of 29 immune cells in TME. Additionally ESTIMATE algorithm was used to estimate the content of Stromal and Immune cells in malignant tumors, and to infer tumor purity and calculate immune score and stromal score.

### Chemotherapy Drug Sensitivity Analysis

The half maximal inhibitory concentration (IC50) of chemotherapy drugs in each sample from TCGA database was estimated via Genomics of Drug Sensitivity in Cancer (GDSC;http://www.cancerrxgene.org/) utilizing the oncoPredict package in R. The differences in drug sensitivity of samples between two risk group were analyzed by Wilcoxon test.

### Statistical Analysis

The patients with CRC were divided into high- and low-risk groups according to the optimal cutoff value. The Kaplan–Meier method was used to evaluate the OS between the high- and the low-risk group, and the log-rank was used to verify the significant difference. The unpaired u-test was used to analyze the distribution of immune cells in the different risk groups. Independent prognostic factors were calculated by Cox proportional hazard regression signature. All statistical analyses were presented via R 3.6.0 (https://www.r-project.org/)., and p < 0.05 was considered statistically different.

## RESULTS

### Patients’ data preparation

491 samples’ of CRC and 42 healthy samples’ RNA-sequencing expression profiles and clinical information were publicly available and downloaded from TCGA-COAD and TCGA-READ cohorts of the TCGA database.

### Identification differentially expressed HSF family genes in normal and tumor samples

Given the close association between HSF family genes and the occurrence and progression of CRC, we performed a differential analysis of HSF family genes in both normal and tumor samples. Our analysis identified HSF4 as the gene exhibiting the most significant difference based on the adjusted P-value. This finding suggests that HSF4 may play a critical role in the pathogenesis of CRC, offering valuable insights into the underlying mechanisms involved in the development of this malignancy.

**Fig. 1.**
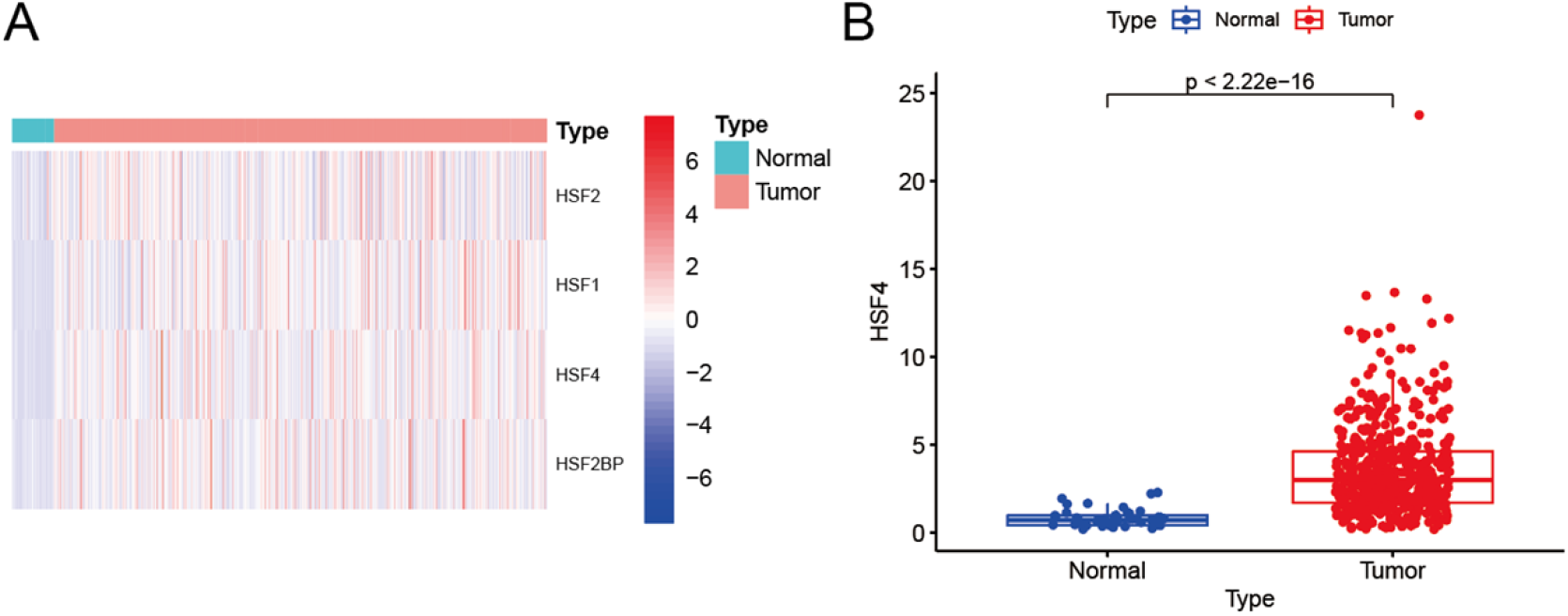
The differential expression of HSF family genes in normal and tumour samples was investigated.

### HSF4’s expression is associated with clinical staging

The expression of certain crucial genes can significantly influence clinical outcomes and tumor staging in patients. Consequently, we investigated the correlation between t HSF4 expression levels and clinical characteristics. Our result revealed a significant association between HSF4 expression and various clinical staging parameters, including stage, T staging, M staging, and N staging. Higher levels of HSF4 expression were correlated with more advanced stages of the disease, suggesting that this gene may contribute to tumor progression. Additionally, we observed Similarly, we found significant associations between HSF4 expression and T, M, and N staging, which are essential indicators of tumor size, metastasis, and lymph node involvement, respectively.

These findings suggest that the expression level of HSF4 could be a potential biomarker for predicting the clinical outcome and staging of patients with certain types of cancers.

**Fig. 2.**
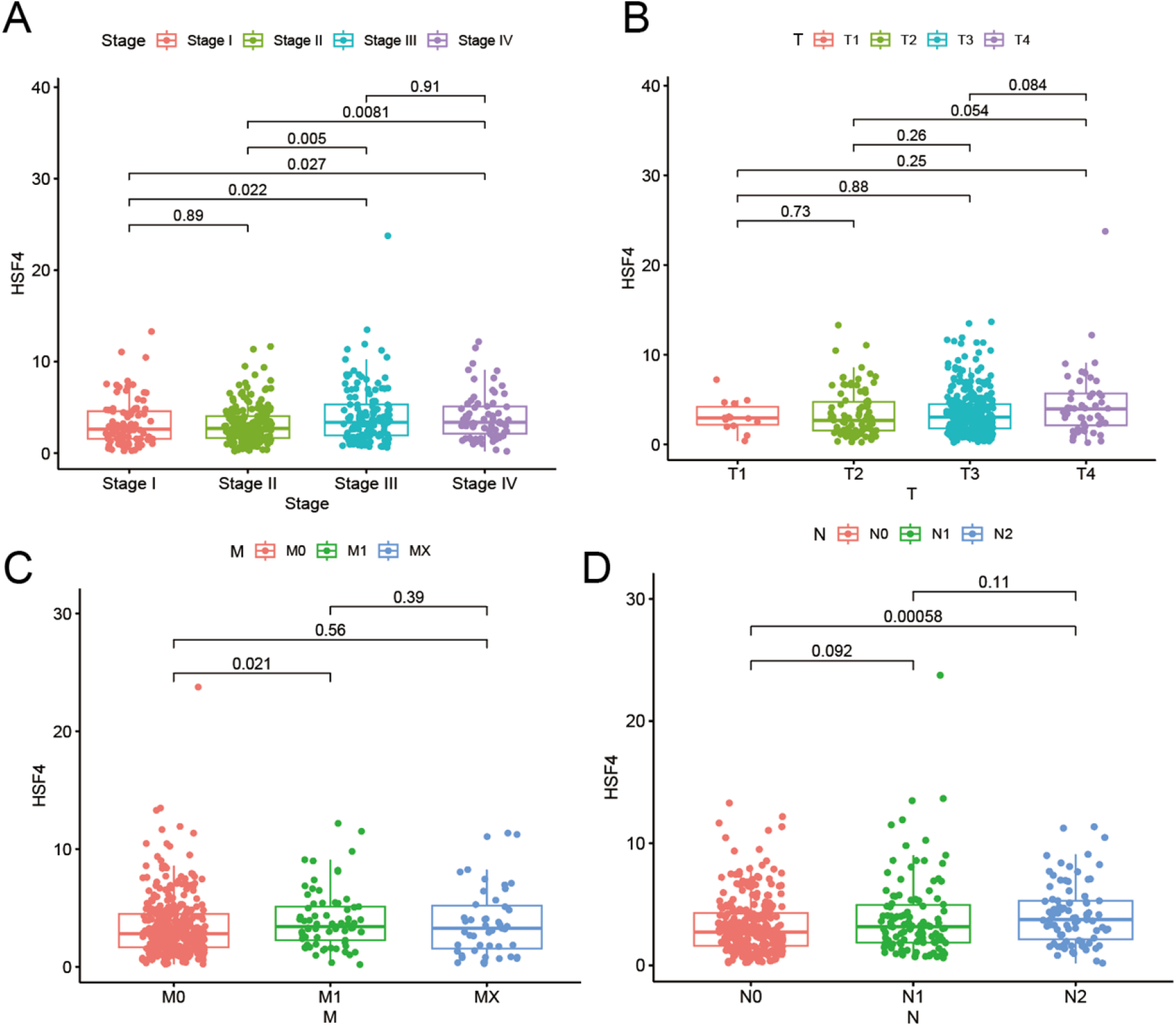
Relationship between clinical staging and HSF4 expression

### HSF4’s expression has a significant impact on prognosis

To investigate the impact of HSF4 expression on the prognosis of CRC patients, we conducted univariate and multivariate Cox regression analysies. The results indicated that HSF4 expression significantly influenced prognosis. This finding suggests that the expression level of HSF4 may be crucial in determining clinical outcomes for patients, potentially offering valuable insights for prognostic prediction and personalized treatment strategies.

### Constructing a risk signature with HSF4’s expression

Based on the above research results, we classified the patients into high- and low-expression groups (high- and low-risk group) using the median value of HSF4’s expression as the threshold. Kaplan–Meier survival indicated that lower expression was associated with better OS (p = 0.047).

**Fig. 3.**
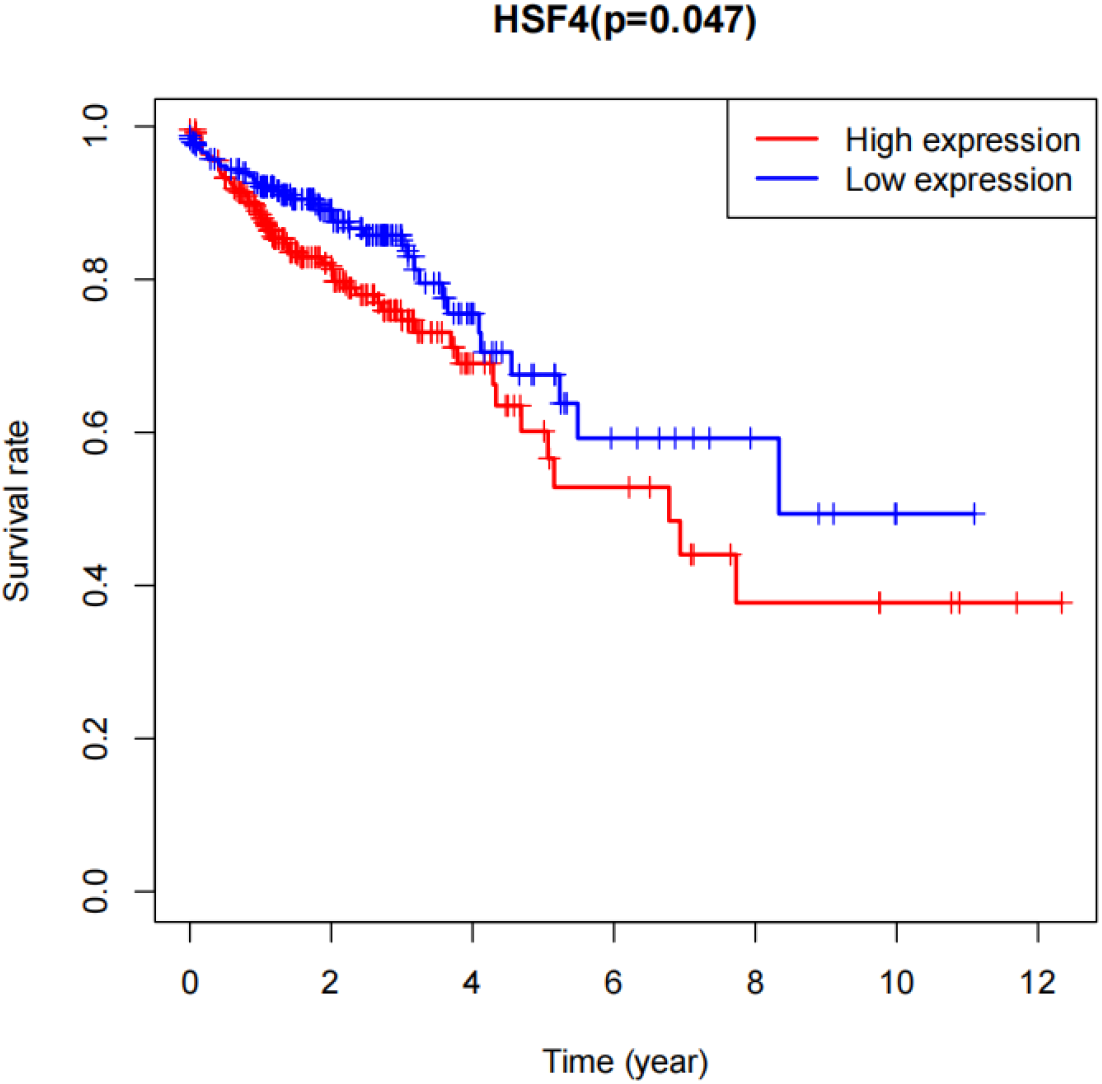
Survival prognosis andHSF4 expression

### Biological pathways of the HSF4-related risk signature

We further focused on exploring the potential mechanisms underlying this phenomenon. To comprehensively analyze the molecular biological characteristics of the HSF4-related risk signature, we identified genes that were strongly associated with it. Using limma software, we generated a heatmap and volcano plot, which revealed 2,494 DEGs with |logFC| > 1 and an adjusted P value < 0.05. Subsequently, we performed GO and KEGG functional enrichment analyses to investigate the underlying mechanisms in greater depth. The GO enrichment analysis indicated that, within the biological process category, these DEGs were primarily involved in cellular responses to type II interferon, Golgi vesicle transport, and protein localization to the cilium. In terms of cellular components, the DEGs were predominantly associated with coated vesicles, nuclear specks, and the coated vesicle membrane. Following this, we conducted KEGG pathway analysis, which revealed that the HSF4-expressed signatures were implicated in several pathways, including endocytosis, human cytomegalovirus infection, and lipid metabolism and atherosclerosis.

**Fig. 4.**
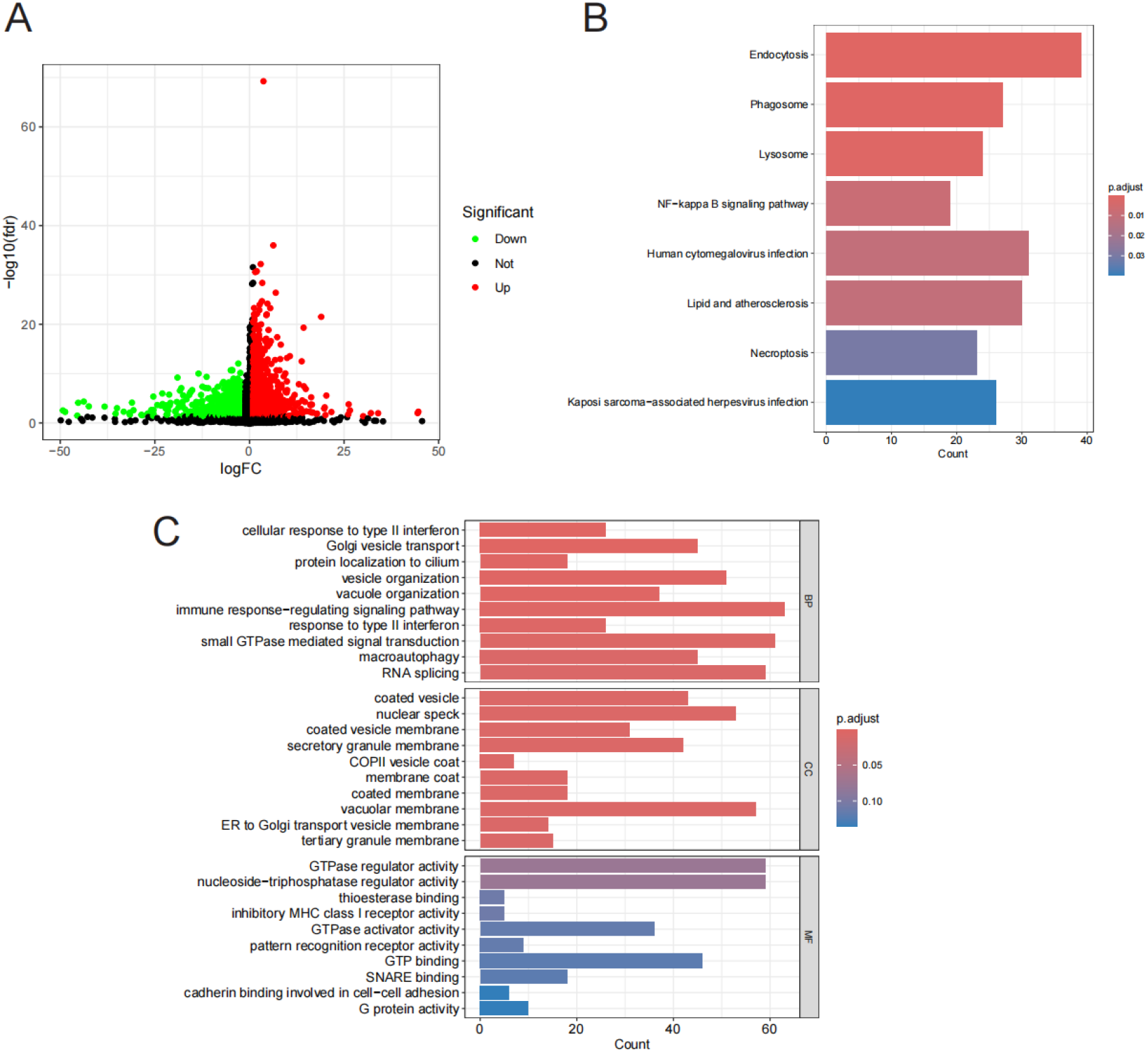
Risk profiles associated with HSF4 and their enrichment results

### HSF4’s expression is positively related to cell proliferation

We conducted a correlation analysis between HSF4 expression and the cell proliferation score,which was calculated using ssGSEA. The results indicated a negative correlation between HSF4 expression and the cell proliferation score (Figure 5B). This negative correlation may reflect the critical role of HSF4 in regulating cell growth and division. HSF4 could serve as a key regulatory factor influencing processes such as the cell cycle, DNA replication, and other mechanisms associated with cell proliferation. Moreover, this negative correlation might also be linked to specific diseases or pathological conditions. In particular, CRC, the expression level of HSF4 mayfluctuate, thereby impacting the rate of cell proliferation and disease progression. These findings suggest that HSF4 has the potential to be a biomarker for therapeutic intervention. Consequently, investigating the relationship between HSF4 and cell proliferation is essential for elucidating the pathogenesis of these diseases and identifying potential treatment strategies.

**Fig. 5.**
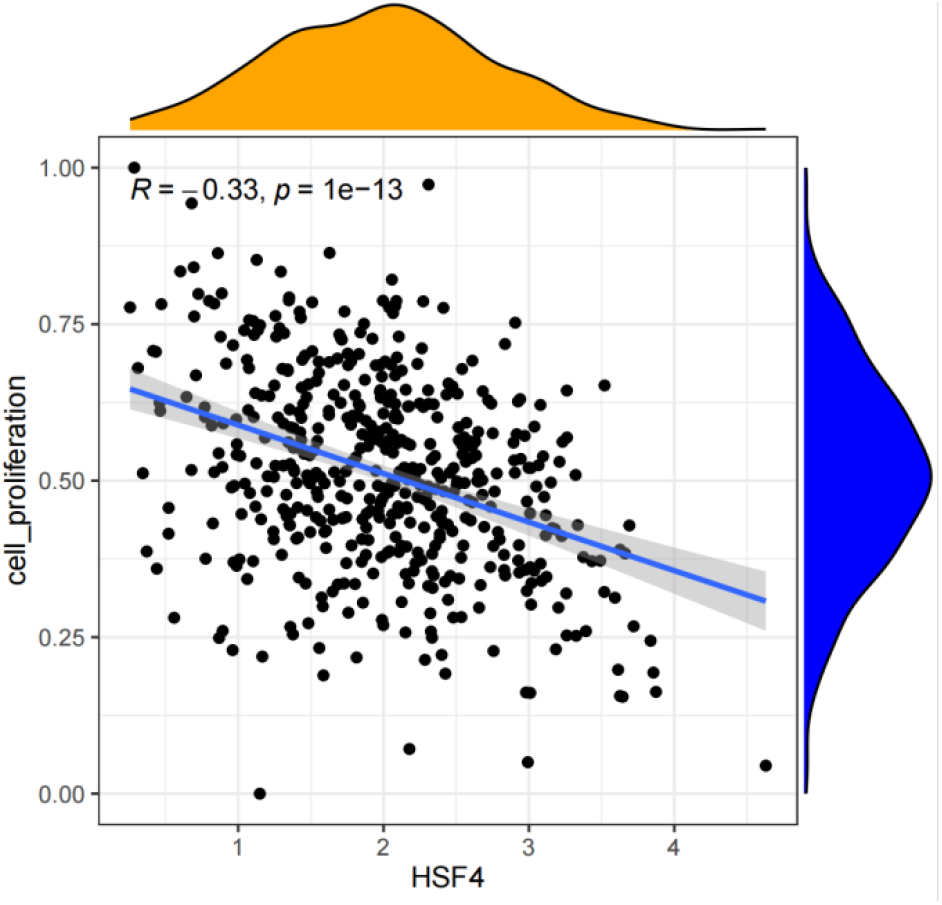
Relationship between HSF4 expression and cell proliferation fraction

**Fig. 6.**
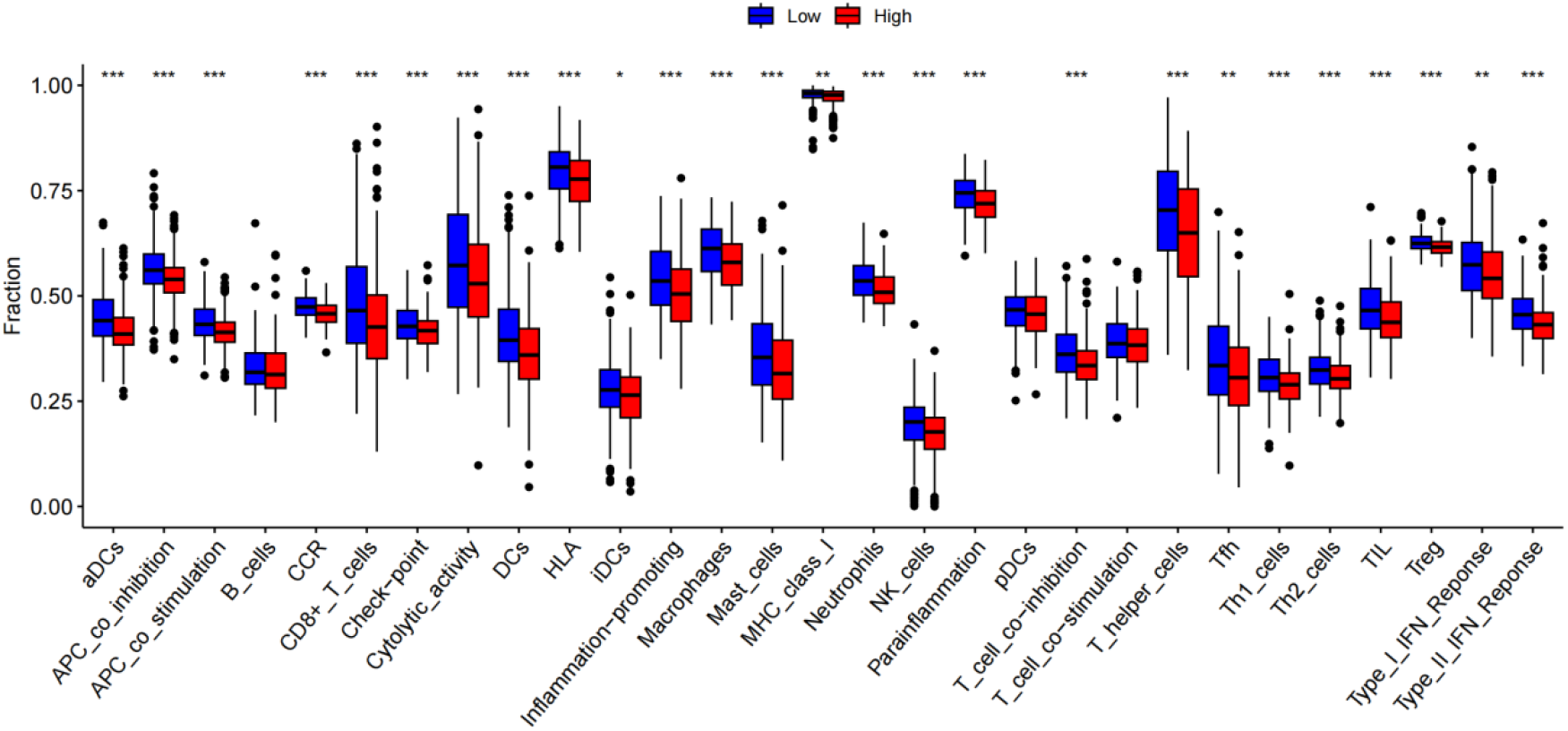
Relationship between HSF4 and immune cell infiltration

**Fig. 7.**
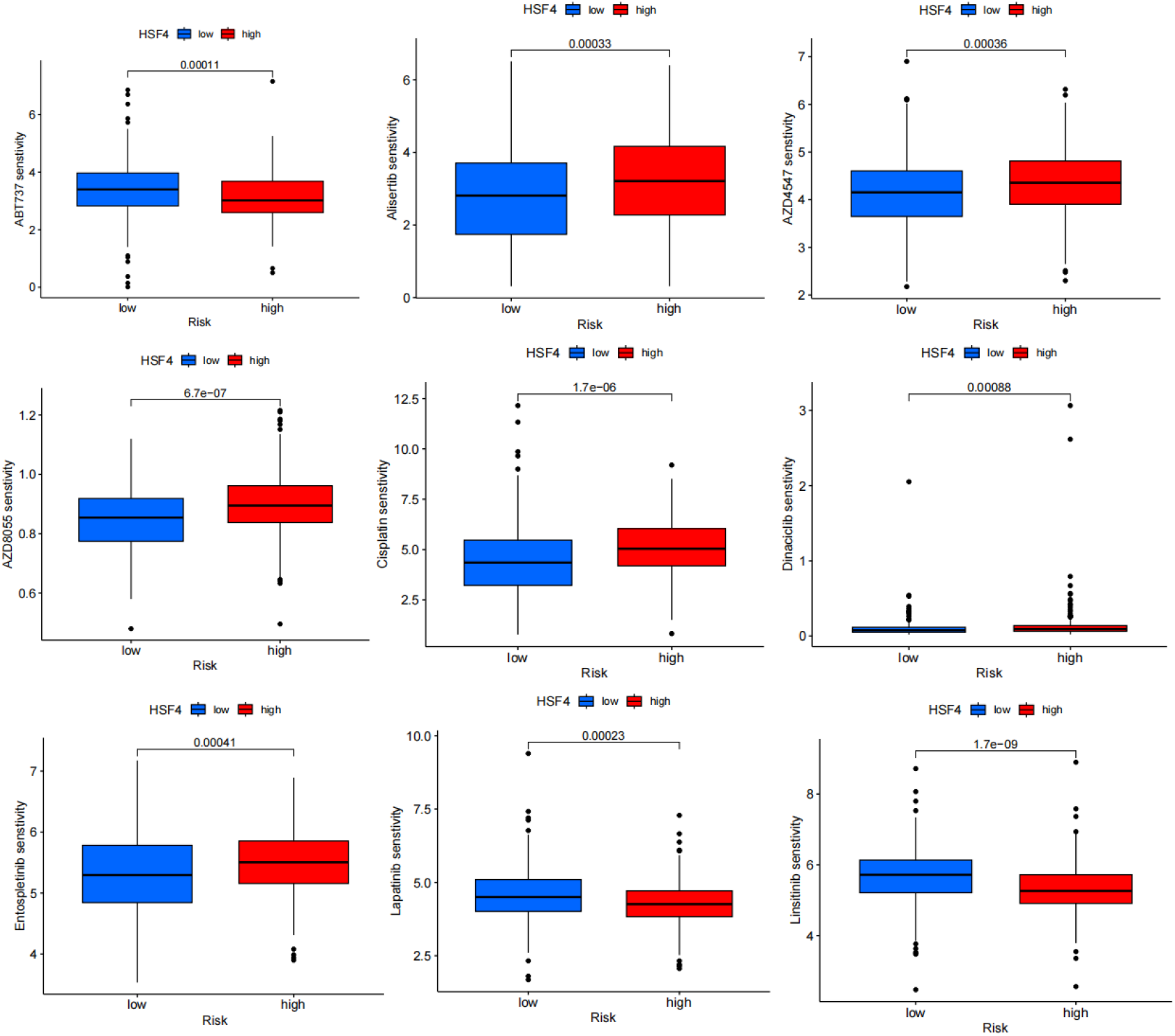
Differences in sensitivity of chemotherapeutic agents between high and low HSF4 subgroups

### Alteration of tumor microenvironment associated with the HSF4-related risk signature

To investigate the relationship between the HSF4-related risk signature and the tumor microenvironment (TME), we employed ssGSEA to assess the differences in 29 types of tumor-infiltrating immune cells between the low-risk and high-risk groups. The results indicated that B cells, pDCs and T helper cells were significantly more enriched in the high-risk group compared to the low-risk group. In contrast, aDCs, CD8+T cells, macrophages, NK cells, DCs, neutrophils and mast cells were predominantly found in the low-risk group.

Additionally, we applied the ESTIMATE algorithm to calculate the estimated score, immune score, stromal score, representing the tumor environment. We found these scores were significantly increased HSF4-low group.

### Evaluation of the Sensitivity of Chemotherapy Drugs to high- and low-risk groups

We further investigated the analysis of drug sensitivity in high- and low-risk groups. Utilizing the GDSC database and the oncoPredict package we conducted a drug sensitivity analysis for each sample, examining the differences in drug sensitivity between the high- and low-expression groups. Ultimately, we identified significant differences in 30 drugs between these groups, suggesting that these 30 agents are potential anticancer drugs for CRC.

## DISCUSSION

The findings from our study provide compelling evidence that HSF4 plays a significant role in the pathogenesis and progression of colorectal cancer (CRC). Differential gene expression analysis revealed that HSF4 is notably upregulated in tumor samples compared to normal tissue, underscoring its potential involvement in cancer development. Furthermore, the correlation between HSF4 expression and advanced clinical stages as well as poor prognosis highlights itscritical role in CRC progression.

The significant correlation between HSF4 expression and key staging parameters, including stage, T, M, and N classifications, indicates that HSF4 expression may directly influence tumor size, metastasis, and lymph node involvement. These associations position HSF4 as a promising candidate biomarker for CRC staging and progression, potentially serving as a valuable tool for clinical assessment and decision-making.

The use of HSF4 expression to stratify patients into high- and low-risk groups has revealed that lower HSF4 expression correlates with better overall survival (OS). This classification into risk groups based on median expression levels not only reinforces the prognostic value of HSF4 but also enhances our understanding of the differential outcomes in patient survival, thereby emphasizing the significance importance of HSF4 in prognostic evaluations.

At the molecular level, the enriched biological pathways associated with HSF4-related differential gene expression, including the response to type II interferon and endocytosis, suggest that HSF4 may influence CRC through immune modulation and cellular trafficking. This may elucidate how variations in HSF4 expression affect cancer cell behaviour and interaction with the tumor microenvironment.

Interestingly, our analysis revealed a negative correlation between HSF4 expression and cell proliferation scores. This counterintuitive finding suggests that HSF4 may play a complex role in cell growth, potentially inhibiting proliferation when overexpressed. Understanding this aspect of HSF4’s functionality is critical for elucidating its dualistic roles in cancer biology, where its effects may vary across different contexts or stages of cancer. Furthermore, the alteration in tumour microenvironment (TME) profiles between high- and low-risk groups based on HSF4 expression was significant. The high-risk group demonstrated an enrichment of B cells, pDCs, and T helper cells, which are typically associated with a more active immune environment. Conversely, the low-risk group displayed higher levels of aDCs, CD8+ T cells, and other immune cells that may contribute to a more effective antitumor response. These findings suggest that HSF4 expression levels could be modulating the immune landscape of CRC, thereby influencing tumor progression and patient outcomes.

The evaluation of chemotherapy drug sensitivity revealed significant differences in response between the high-risk and low-risk groups. This finding underscores the potential of utilizing HSF4 expression levels not only as a biomarker for disease progression but also for the customization of chemotherapy treatments. The identification of 30 drugs with varying sensitivities between these groups establishes a foundation for personalised treatment approaches,which may enhance therapeutic efficacy and improve patient survival.

## CONCLUSION

In conclusion, our comprehensive analysis of the HSF family genes in colorectal cancer (CRC) has revealed HSF4 as a gene exhibiting significant differential expression between normal and tumor tissues. We further established its prognostic significance through both univariate and multivariate Cox regression analysis. The expression level of HSF4 was significantly associated with clinical characteristics including tumor stage, T stage, M stage, and N stage. Our analysis also demonstrated a positive correlation between HSF4 expression and cell proliferation, suggesting its potential role as a therapeutic target. Additionally, we evaluated the tumor microenvironment and immune cell infiltration, which provided insights into the tumor immune landscape. Finally, drug sensitivity analysis revealed 30 potential anti-CRC drugs that exhibited significant differences in sensitivity between the high and low HSF4 expression groups. Collectively, our findings underscore the critical role of HSF4 in CRC and propose its potential as both a prognostic marker and therapeutic target. These results offer valuable insights for future research on CRC pathogenesis and the development of personalized treatment strategies.

## SUPPLEMENTARY MATERIAL

**Supplementary. 1.**
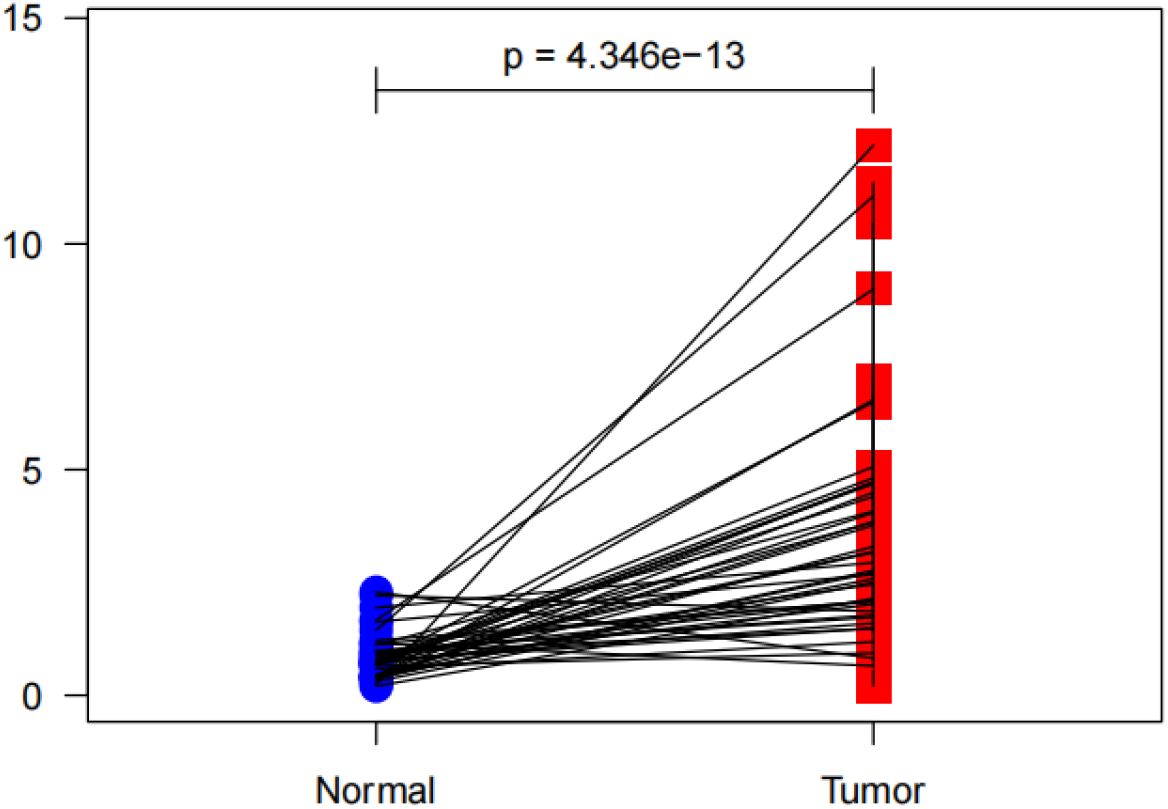
Differential expression of HSF4 between paired subgroups

**Supplementary. 2.**
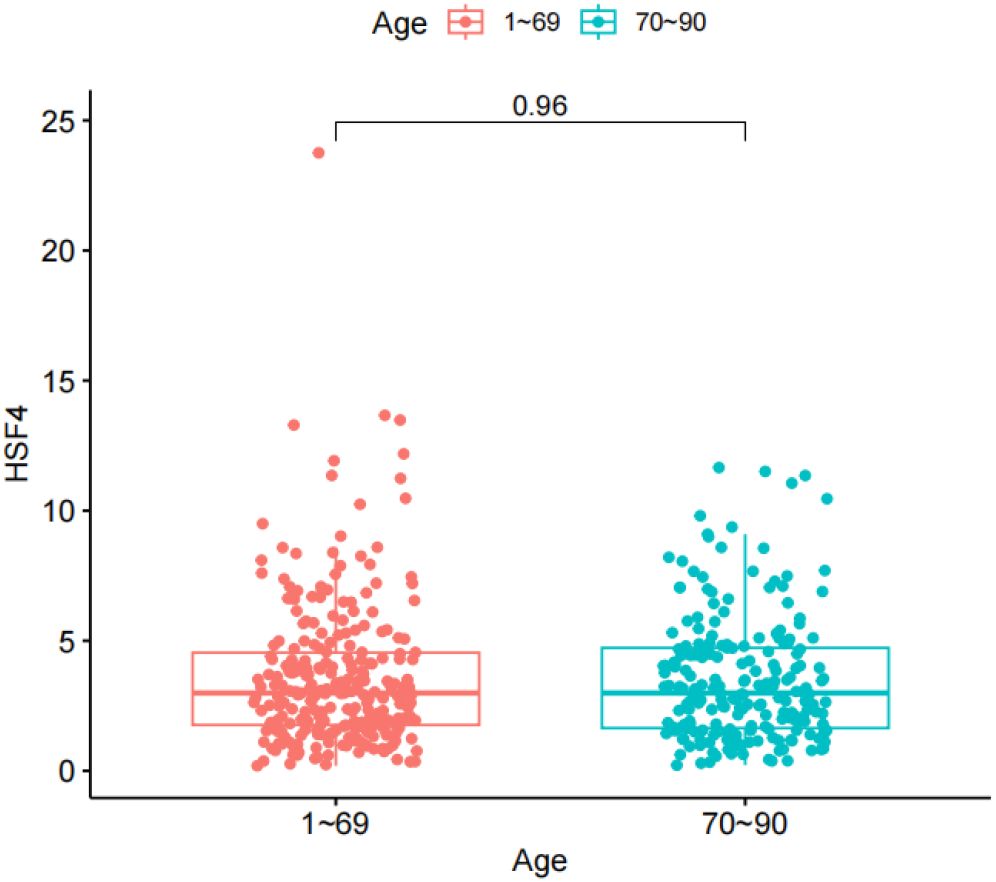
Differences in HSF4 expression between age subgroups

**Supplementary. 3.** Logistic regression relationships between HSF4 expression and clinical information.

| Clinical Characteristics | Total | Odds ratio in HSF4 expression | P-Value |
| --- | --- | --- | --- |
| Stage ( I VS IV) | 158 | 1.138889 (0.6825523-1.91271) | 0.01098282 |
| Age (<70 VS ≥70) | 488 | 1 (0.6996522-1.429282) | 1 |
| T (T1 VS T4) | 65 | 0.9934641 (0.3190016-3.242363) | 0.2911443 |
| M (M0 VS M1) | 434 | 1.876293 (1.113143-3.216303) | 0.01961575 |
| N (N0 VS N3) | 373 | 1.386736 (0.8984372-2.14528) | 0.001119075 |

**Supplementary. 4.**
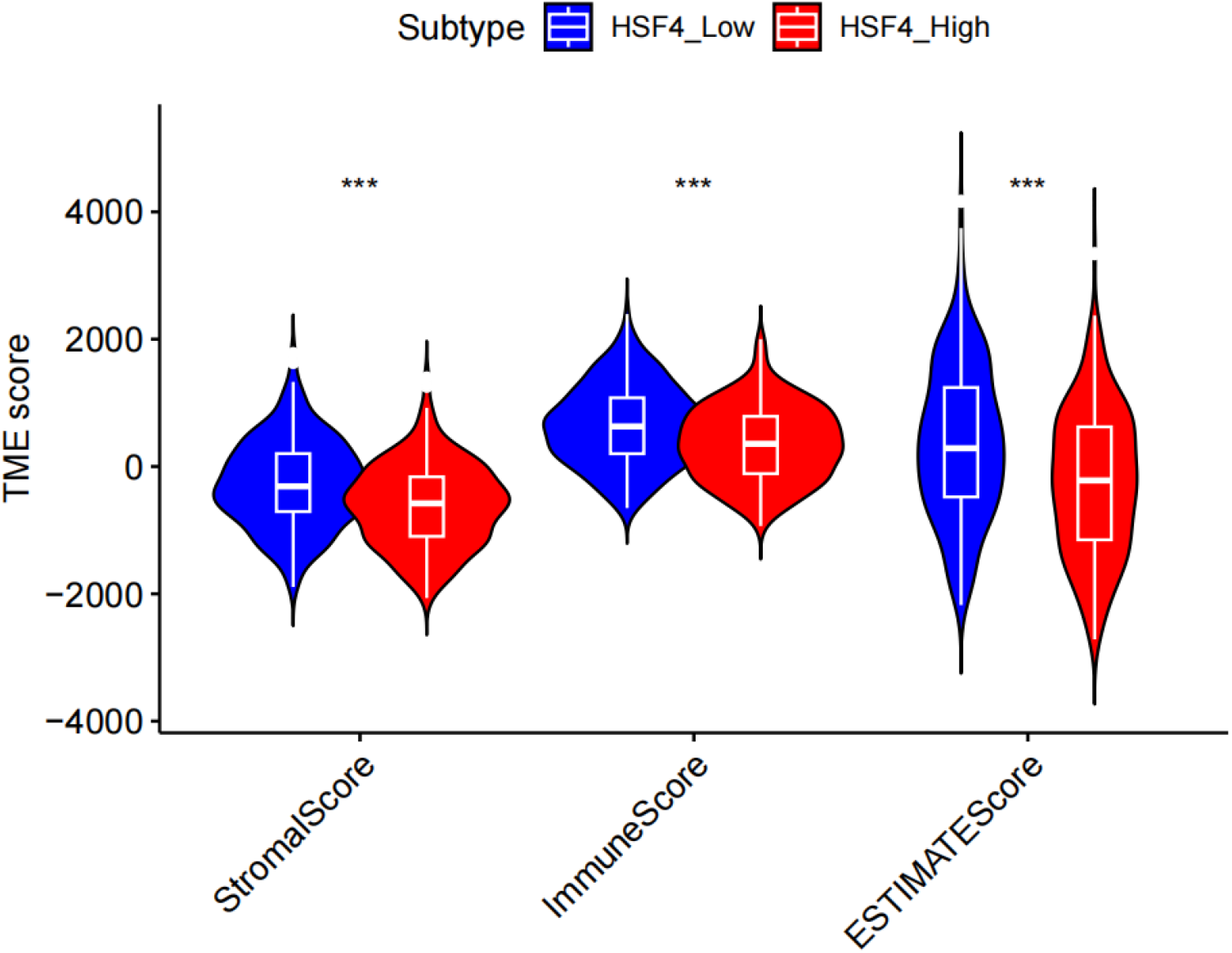
Differences in TME scores between groups with high and low HSF4 expression

